# Eyes closed or Eyes open? Exploring the alpha desynchronization hypothesis in resting state functional connectivity networks with intracranial EEG

**DOI:** 10.1101/118174

**Authors:** Jaime Gómez-Ramírez, Shelagh Freedman, Diego Mateos, José Luis Pérez-Velázquez, Taufik Valiante

## Abstract

This paper addresses a fundamental question, are eyes closed and eyes open resting states equivalent baseline conditions, or do they have consistently different electrophysiological signatures? We compare the functional connectivity patterns in an eyes closed resting state with an eyes open resting state, and show that functional connectivity in the alpha band decreases in the eyes open condition compared to eyes closed. This "alpha desynchronization " or reduction in the number of connections from eyes closed to eyes open, is here, for the first time, studied with intracranial recordings. We provide two calculations of the wiring cost, local and mesoscopic, defined in terms of the distance between the electrodes and the likelihood that they are functionally connected. We find that, in agreement with the "alpha desynchronization" hypothesis, the local wiring cost decreases going from eyes closed to eyes open. However, when the wiring cost calculation takes into account the connectivity pattern, the wiring cost variation from eyes closed to eyes open is not as consistent and shows regional specificity. The wiring cost measure defined here, provides a new avenue for understanding the electrophysiology of resting state.

## 1 Introduction

The view of the brain as a reflexive organ whose neural activity is completely determined by incoming stimuli is being challenged by the "intrinsic" or spontaneous view of the brain. Nevertheless, the exact implications of resting state for brain function are far from clear (Schneider et al., 2008), (Northoff et al., 2010). Mandag and colleagues (Maandag et al., 2007) argue for the reconceptualization of resting state as an independent variable (brain's input) to a multidimensional activity modulator. The emerging field of functional connectomics relies on the analysis of spontaneous brain signal covariation to infer the spatial fingerprint of the brain's large-scale functional networks. While there is growing interest in the brain's resting state, supported by evidence for persistent activity patterns in the absence of stimulus-induced activity (e.g. default mode network (Greicius and Menon, 2004)), there is not a definite recommendation about whether resting state data should be collected with participants' eyes open or closed. If stimulus-induced activity is indeed, at least in part, predetermined by the brain's intrinsic activity (i.e. resting state activity), it follows that we cannot understand one without the other. The more we know about the electrophysiological underpinnings of resting state, both with eyes closed and eyes open, the better equipped we will be to understand brain dynamics, including both intrinsic activity and the processing of stimuli.

The orthodox approach to understanding brain function relies on the view of the brain as an organ that produces responses triggered by incoming stimuli, which are delivered at will by an external observer. This idea has been challenged by the complementary view of the brain as an active organ with intrinsic or spontaneous activity (Llinás, 1988); (Biswal et al., 1995); (Papo, 2013). Crucially, the brain’s intrinsic activity both shapes and is shaped by external stimuli. While there has been some controversy concerning the ecological relevance of studying a default or resting condition (Buckner and Vincent, 2007); (Morcom and Fletcher, 2007), the empirical evidence for intrinsic brain activity is conclusive (Wang et al., 2006); (Mantini et al., 2007).

Despite the ever increasing importance of resting-state functional connectivity (a quick search on PubMed shows 2,742 papers with the term "resting state" in the title at the time of the writing), it remains underutilized in clinical decision making (Tracy and Doucet, 2015). A rationale for this needs to be found in both conceptual and methodological basis. First and foremost, the term resting-state is a misnomer, as a matter of fact, the brain is always active, even in the absence of an explicit task. Cognitive task-related changes in brain metabolism measured with PET account for only 5% or less of the brain’s metabolic demand (Sokoloff et al., 1955). Second, the resting state literature from its inception is eminently based on the analysis of low frequency fluctuations of the BOLD signal measured using fMRI, alone or in combination with EEG and PET (Van Den Heuvel and Pol, 2010); (Musso et al., 2010). Third, these techniques suffer from suboptimal temporal and/or spatial resolution and the haemodynamic or metabolic activity measured in fMRI and PET are proxy measures for the electrophysiological activity. Fourth, there is a lack of consensus in the literature regarding whether resting state data should be collected while the participant has their eyes open, closed or fixated. See (Patriat et al., 2013) for non-significant between-condition differences in resting state networks and (Yan et al., 2009) for an antagonistic view. This paper attempts to better understand the brain's resting state by characterizing the two most common baseline conditions in neuropsychology, eyes closed and eyes open, using intracranial electroencephalogram (iEEG).1

Previous studies have identified a reduction in the number of connections when the closed eyes condition is compared to the eyes open condition, in the alpha band (Tan et al., 2013); (Barry et al., 2007).

This is known as "alpha desynchronization". Using EEG, Barry and colleagues (Barry et al., 2007)found that there are electrophysiological differences-topography as well as power levels-between the eyes closed and eyes open resting states. A higher degree of alertness caused by opening one's eyes is associated with the attenuation of alpha rhythm, which is supplanted by desynchronized low voltage activity (Niedermeyer and da Silva, 2005). Geller and colleagues (Geller et al., 2014)found that eye closure causes a widespread low-frequency power increase and focal gamma attenuation in the human electrocorticogram. However, although these studies explicitly make the case that eyes open and eyes closed are different baseline conditions, they do not provide a method for comparing the functional connectivity patterns elicited by either of the two conditions against a common criterion.

Shedding some light on the problem, this paper examines whether the eyes closed and eyes open resting states are equivalent baseline conditions by analyzing the differences between the two. First, we perform power and phase based connectivity analysis to asses whether the connectivity patterns calculated from intracranial recordings are able to differentiate between the two conditions. Second, we calculate the wiring cost for the connectivity maps (in two ways, local and global) for each condition, to investigate whether the wiring cost can be used as a feature/covariate to distinguish between the eyes closed and eyes open conditions.

## 2 Materials and Methods

### 2.1 Participants

The intracranial electroencephalography recordings were collected at the Toronto Western Hospital (Toronto ON, Canada). Our research protocol was approved by the institutional review board and informed consent was obtained from the participants. 11 participants (6 female) with pharmacologically-refractory mesial temporal lobe epilepsy underwent a surgical procedure, in which electrodes were implanted sub-durally on the temporal lobe and stereotaxic depth electrodes located in the hippocampi or other deep structures (Figure 1). For each patient, electrode placement was determined to best pinpoint the origin of seizure activity. In addition to electrodes implanted in the temporal lobe, including depth electrodes in the hippocampi, some patients had electrodes implanted in frontal, interhemispheral and the cortical convexity (see Table 1). The electrode implants are thus not identical for all participants, though they tend to overlap in the mesial temporal lobe epilepsy (MTLE) sensitive regions. This limits the ability to directly compare the wiring cost or other network properties among participants. However, we can still compare and generalize participants by examining the difference between the two conditions. For example, in order to compare the functional connectivity pattern for two participants, one with a grid in the left cortex and another with depth electrodes in the hippocampus and temporal areas, we calculate the difference between network parameters from eyes closed to eyes open, within each participant.

**Figure 1:**
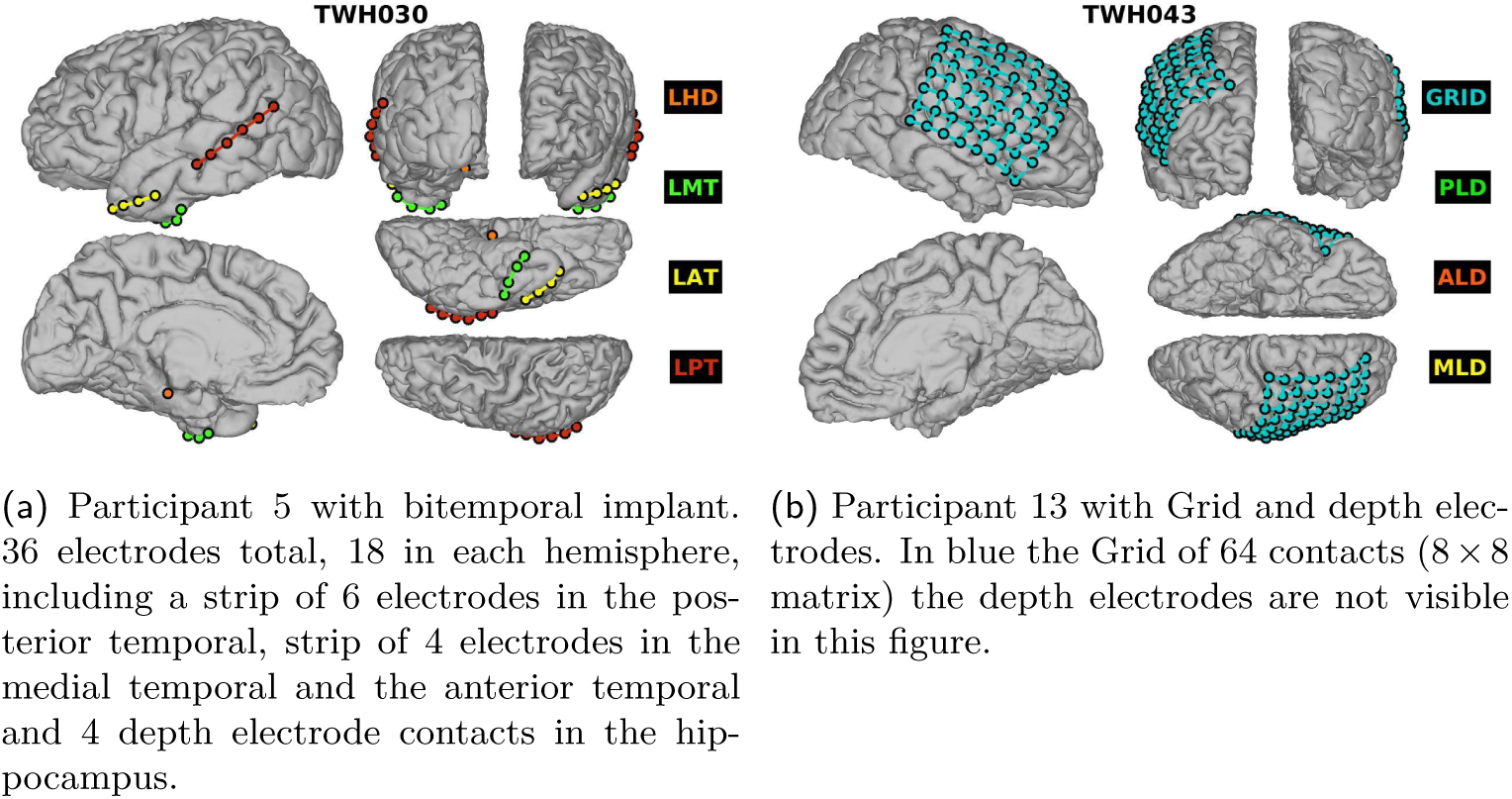
Schematic of the electrode implant for two participants

**Table 1:** ID, sex, age, laterality and type of implant. The laterality can be bilateral (B), left (L) and right (R). The location of the electrodes fall under the following categories: hippocampus (H), temporal (T), frontal (F), interhemispheral (IH), frontal polar (FP), Grid, and depth (D; different from hippocampus).

| Patient | Sex | Age | Laterality | Channels |
| --- | --- | --- | --- | --- |
| 5 | F |  | B | H, T |
| 6 | M |  | B | H, T, F, IH |
| 7 | M |  | B | H, T, F, IH |
| 10 | F |  | B | FP, IH, F |
| 11 | F |  | B | H, T |
| 12 | M |  | B | H, T |
| 13 | F |  | B | Grid, D |
| 15 | F |  | R | H, FP, F, IH, T |
| 16 | F |  | B | H, T |
| 17 | M |  | L | Grid, D |
| 18 | M |  | R | H, FP, F, T |

### 2.2 Resting state conditions

To assess resting-state activity for both the eyes closed and and eyes open conditions, participants were asked to relax and rest quietly in their hospital bed, in a semi-inclined position. First, they were asked to close their eyes for three minutes and then asked to keep their eyes open for another three minutes. Each session was recorded with real-time monitoring of the intracranial electroencephalography and continuous audio and video surveillance. ECoG recordings allow us to simultaneously study both fast and slow temporal dynamics of the brain at rest, that is, not engaged in tasks prescribed by the experimenter. Freeman and Zhai (Freeman and Zhai, 2009)have shown that the resting ECoG is low-dimensional noise, making resting state an optimal starting point for defining and measuring both artifactual and physiological structures emergent in the activated electrophysiological signals. Importantly, ECoG signals co-vary in patterns that resembled the resting state networks (RSN) found with fMRI (Fukushima et al., 2015).

### 2.3 iEEG acquisition

Continuous iEEG data were recorded in an unshielded hospital room using NATUS Xltech digital video-EEG system. Commercially available, hybrid depth electrodes and subdural electrodes were used to collect continuous iEEG recordings. Common reference and ground electrodes were placed subdurally at a location distant from any recording electrodes with contacts oriented toward the dura. Electrode localization was accomplished by localizing the implanted electrodes on the postoperative computed tomography (CT) scan. Subdural electrodes were arranged in strip configurations (4, 6 or 8 contacts) with an interelectrode spacing of 10 mm. The location of the electrode implants is not identical across patients, however, all participants had depth electrodes, mostly in the hippocampi (Table 1).

### 2.4 Signal processing

Signals were filtered online using a high-pass (0.1 cutoff frequency) and an anti-aliasing low-pass filter. Offline filtering using Matlab in house-scripts, consisted of a high-pass and low-pass filter at 0.5-70 Hz and a notch filter applied at 60 Hz to remove electrical line noise.

To extract power and phase estimates of time-varying frequency-specific, the ECoG signals were convolved with complex-valued Morlet wavelets. The wavelet convolution transformed the voltage trace at each electrode to obtain both instantaneous power and phase trace for each frequency. The wavelet length was defined in the range of −1 to 1 seconds and was centered at *time* = 0 (in doing so we guarantee that the wavelet has an odd number of points). We used a constant number of wavelet cycles (7). This number was chosen since we have long trial periods (3 minutes) in which we expect frequency-band-specific activity, and a large number of cycles (from 7 to 10) facilitates identifying temporally sustained activity (Cohen, 2014).

Of note, it is also possible to use a number of wavelet cycles that changes as a function of frequency, to adjust the balance between temporal and frequency precision as a function of the frequency of the wavelet. Thus, there is a trade-off between temporal and frequency precision. Since we are processing long epochs, we favour frequency over time precision and therefore chose to use a large number of wave cycles.

### 2.5 Connectivity measures

We are interested in calculating the wiring cost associated with the functional connectivity map defined upon the electrodes’ spatial location. Functional connectivity is calculated using both power-based and phase-based measures. For power-based we calculate Spearman’s correlation and we calculate two different measures for phase-based connectivity, phase-lag index (Stam et al., 2007) and intersite phase clustering ISPC^2^

Here we briefly outline the three connectivity measures used. First, we describe power-based connectivity and next phase-based connectivity for the phase lag index (PLI) and intersite phase clustering (ISPC) measures. The last part of this section describes the computation of the wiring cost associated with the functional connectivity map.

#### 2.5.1 Power-based connectivity

To calculate the correlation coefficients for power time series from any two electrodes in the same frequency, we perform time-frequency decomposition using wavelets to then compute the Spearman correlation coefficient between the power time series of the two electrodes.

To increase the signal to noise ratio, we segment the data into non-overlapping windows of 5 seconds, compute Spearman’s correlation coefficient for each segment, and then average the correlation coefficients together.

The Spearman’s correlation is the Pearson correlation of the data previously rank-transformed. Formally, the Spearman correlation of two channels *x* and *y* whose power time series values have been rank-transformed is:

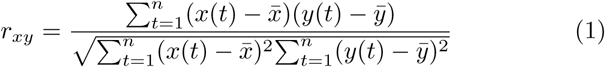

It is of note that power-based correlation coefficients range from −1 and 1. To have a more normal looking distribution, it is preferable to perform a Fisher-Z transformation.

#### 2.5.2 Phase-based connectivity

We calculate phase-based connectivity with two different measures, intersite phase clustering (ISPC) and the phase-lag index (PLI). The ISPC measures the clustering in polar space of phase angle differences between electrodes and is given by the equation:

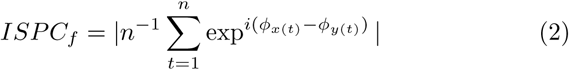

where n is the number of time points and *ϕ_x_* and *ϕ_y_*are the phase angles from electrodes *x* and *y* at a given frequency *f*. Note that this measure can be sensitive to volume conduction. For example, when the phase differences are not uniformly distributed, but clustered around 0 or *π* in polar space, much of the apparent connectivity between these electrodes might be due to volume conduction.

There are several phase-based connectivity measures that ignore the 0 – *π* phase-lag connectivity problem, e.g., imaginary coherence (Nolte et al., 2004), phase-slope index (Nolte et al., 2008), phase-lag index (Stam et al., 2007) and weighted phase-lag index Vinck et al. (2011). Although these measures are designed to be insensitive to volume conduction, in some cases they may still be susceptible to mixing sources (Peraza et al., 2012).

Phase lag index (PLI) measures the extent to which the distribution of phase angle differences is more to the positive or to the negative side of the imaginary axis on the complex plane. That is, it tells us whether the vector of phase angle differences are pointing up or down in polar space. The idea is that if spurious connectivity is due to volume conduction, the phase angle differences will be distributed around zero radians. It follows that non-volume conducted connectivity will produce a distribution of phase angles that is predominantly on either the positive or the negative side of the imaginary axis. Note that here, contrary to ISPC, the vectors are not averaged, instead is the sign of the imaginary part of the cross spectral density that is averaged.

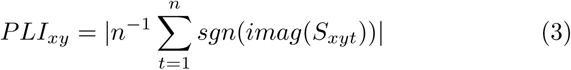

where *imag* is the imaginary part of *S_xy_*_(t)_ or cross-spectral density between channels *x* and *y* at time *t*. The *sgn* function returns +1,−1 or 0.

To recapitulate, ISPC captures the clustering of the phase angle difference distribution and PLI the phase angle directions. ISPC can be influenced by changes in power and is maximally sensitive to detecting connectivity, regardless of the phase angle differences.

Again, we calculate phase-based connectivity with two different measures, ISPC and PLI. Phase coherence measures are highly influenced by volume conduction (Mormann et al., 2000). PLI, on the other hand, was designed to tackle this problem Stam et al. (2007). Peraza and colleagues (Peraza et al., 2012)show that PLI is partially invariant to volume conduction. In a simulation study they found that PLI-based connectivity networks show more small worldness (higher cluster coefficient) than random networks. But, for non-volume conduction, PLI-based networks are close to random networks, indicating that the high clustering shown for PLI is caused by volume conduction. Therefore, PLI is not insensitive to volume conduction. Intracranial EEG is less sensitive to volume conduction problems than other electrophysiological techniques (EEG and MEG). Thus, by calculating phase-based connectivity with both ISPC and PLI, we expect to clarify the properties of both measures for the analysis of the iEEG signal.

#### 2.5.3 Wiring cost

Now that we have described how functional connectivity is obtained, we continue describing how to calculate the wiring cost between any pair of electrodes. To begin, we define the distance matrix *D_ij_ =||*(*x_i_,y_i_,z_i_*),(*x_j_, y_j_, z_j_*)|| which captures the Euclidean distance between any two electrodes physically located in Cartesian coordinates (*x_i_, y_i_, z_i_*) and (*x_j_,y_j_,z_j_*). Second, we calculate the functional connectivity for each condition, eyes closed and eyes open, using the connectivity matrices described above. Two Spearman correlation of power time-series and two phase-based connectivity criteria-inter site phase clustering (ISPC) and phase lag index (PLI).

The computation of the wiring cost *W* combines a physical distance matrix *D* and a functional connectivity matrix *F*. While there is one phyisical distance matrix *D* for each participant, the functional connectivity matrix *F* can be calculated using three different criteria -ISPC, PLI and the Spearman correlation of power time series.

Plus, we define two methods for calculating the wiring cost. First, as a measure of the cost of moving information between two electrodes, taking into account only its distance (pairwise wiring cost) and second, taking into account the connectivity pattern, as well as the distance (mesoscopic wiring cost).

The pairwise wiring cost for a distance matrix of electrodes *D* and functional connectivity matrix *F* calculated at frequency *f* is calculated as:

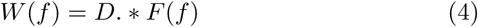

Thus, the pairwise wiring cost of two electrodes is directly proportional to the distance and the correlation. The further away and the stronger the correlation, the larger the wiring cost (Figure 2).

The mesoscopic wiring cost, on the other hand, is directly proportional to the Euclidean distance that separates the electrodes and inversely proportional to the likelihood that the two electrodes are functionally connected. For example, for two electrodes *A* and *B*, the wiring cost is defined as follows

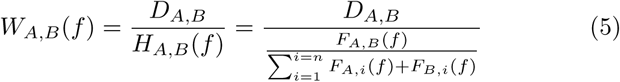

In words, the wiring cost between two electrodes *A, B* in frequency *f* is the Euclidean distance *D_A,B_* divided by the likelihood that *A* and *B* are functionally connected *H_A,B_*, which is calculated as the ratio between the functional connectivity *F_A_,_B_* and the sum of the functional connectivity between *A* and *B* and all their neighbours (See Figure 2). Thus, the mesoscopic wiring cost increases with distance (the further away two points are, the more energy required to move information between them) and decreases with the odds that the two electrodes are functionally connected relative to other possible connections. That is, for the same distance *D_A_,_B_*, the more likely *A* and *B* are connected, the less the wiring cost, or in other words, the more rare the functional connection, the greater the wiring cost. Figure 3 describes the algorithm employed to calculate the wiring cost for each participant.

**Figure 2:**
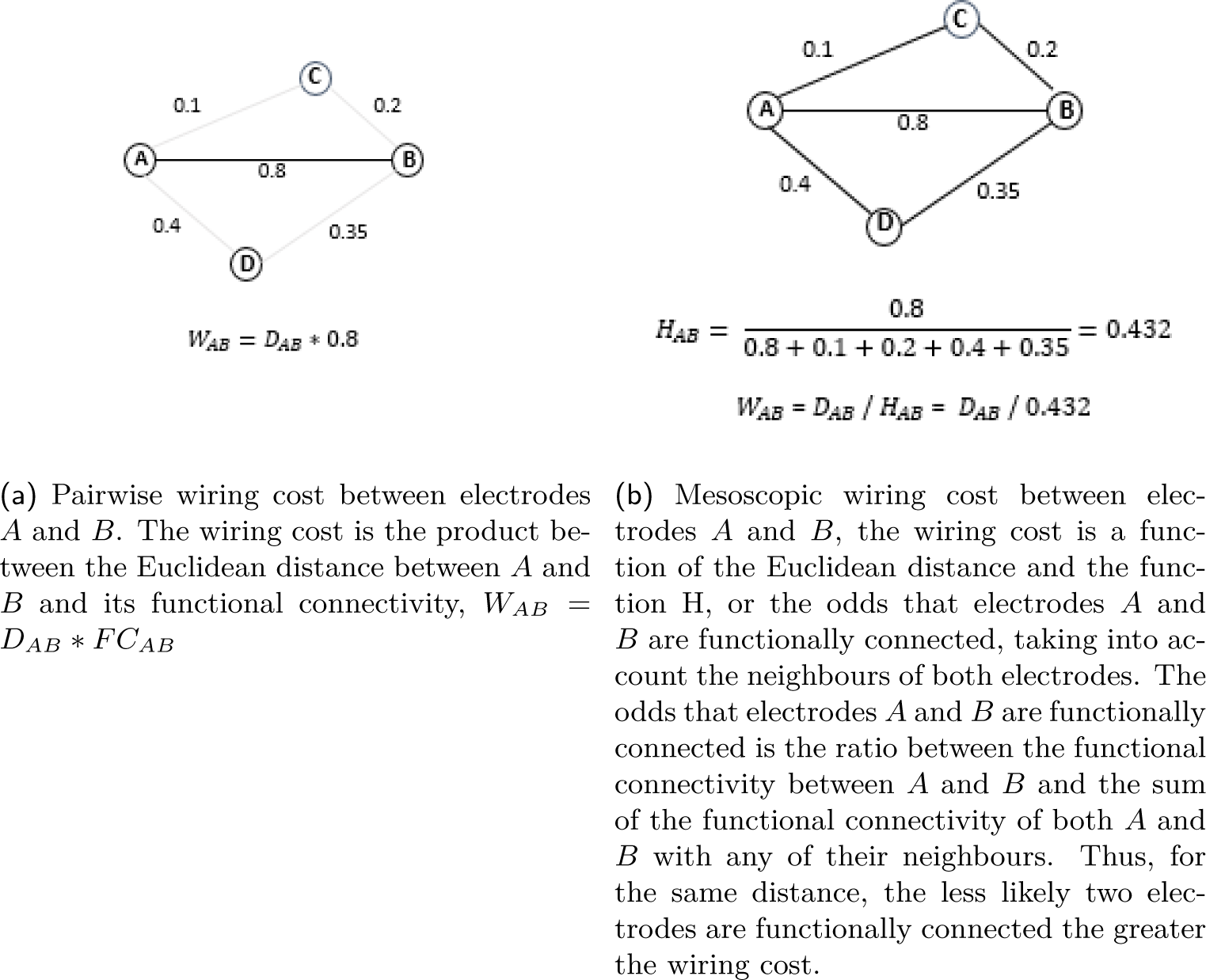
Calculation of the pairwise wiring cost between two electrodes (2a) and the mesoscopic wiring cost between two electrodes, which also takes into account the connectivity pattern of the electrodes (2b)

**Figure 3:**
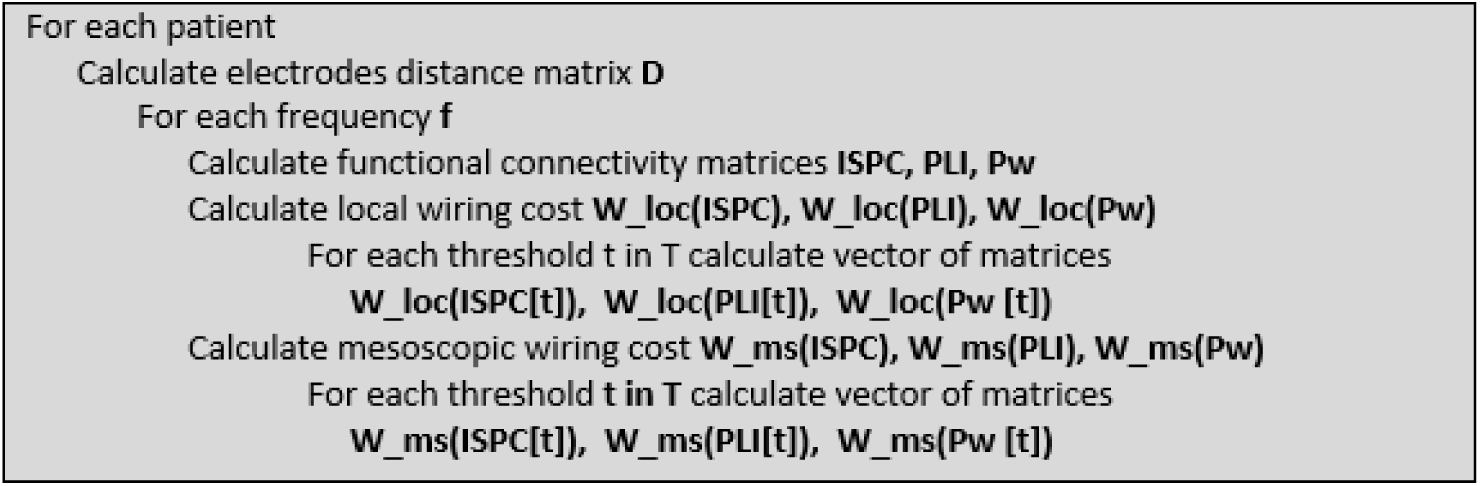
Algorithm used to calculate the wiring cost from the electrodes distance-matrix D and the functional connectivity matrices. Each subject has one matrix D and one connectivity matrix per condition (eyes closed and eyes open) and frequency band in which the functional connectivity is calculated using three different measures, ISPC, PLI and power correlation. Finally, each wiring cost matrix is thresholded to obtain a binary matrix, one for each threshold.

## 3 Results

We systematically explore the electrophysiological underpinnings of resting state, by analyzing both the eyes closed and eyes open conditions with intracranial electroencephalogram data (iEEG). The excellent temporal and spatial precision of ECoG allows us to explore the electrophysiology of these two resting states in an optimal way. We perform connectivity analysis to characterize both baseline conditions, using an energy cost efficiency approach.

We calculate the wiring cost per participant as the average of the wiring cost for each pair of electrodes according to Equations 4 and 5.The Euclidean distance matrix between electrodes and the functional matrices for the power-based and both phase-based connectivity measures, give us three wiring cost matrices (phase-lag index, intersite phase clustering and power correlation) per participant, condition and frequency. Thus, 3 different connectivity measures, 11 patients, 2 conditions and 6 frequencies, yields a total number of wiring cost values equal to 3 × 11 × 2 × 6 = 396. However, we are interested in the difference between eyes closed and eyes open, therefore the wiring cost is a vector half the size, or 3 × 11 × 1 × 6 = 198.

### 3.1 Pairwise wiring cost

We start by calculating the pairwise wiring cost (Equation 4). Figure 4 shows the difference between the pairwise wiring cost for eyes closed and eyes open averaged across participants, with the functional connectivity matrices calculated for 6 frequency bands: delta, theta, alpha, low and high beta, and gamma. The first two plots in Figure 4 show the wiring cost difference for phase-based connectivity -ISPC (Equation 2) and PLI (Equation 3)- and the last plot depicts the wiring cost difference for power-based connectivity (Equation 1).

**Figure 4:**
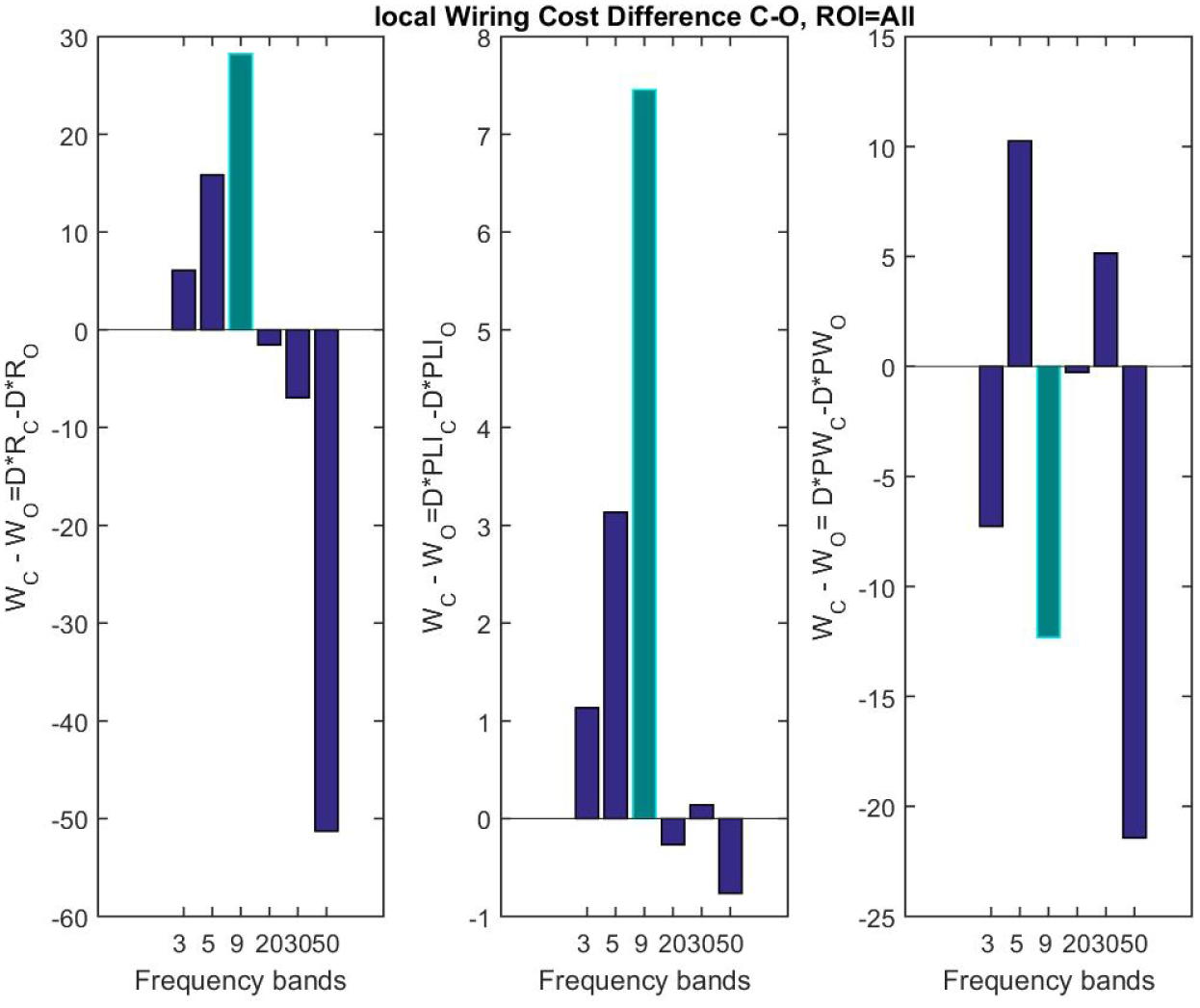
Figure shows the difference in wiring cost between the eyes closed and eyes open conditions, averaged for all patients across six frequency bands, delta, theta, alpha, low and high beta, and gamma. The alpha band is highlighted in green. The wiring cost difference between eyes closed and eyes open is *D*(*F_C_* (*f*) – *F_O_*(*f*)), where *D* is the matrix containing the Euclidean distance between any two electrodes and matrices *F_C_* (*f*) and *F_O_* (*f*) represent the functional connectivity for eyes closed and eyes open respectively, in the frequency band *f*. From left to right, the wiring cost difference for phase-based connectivity using ISPC, PLI and power-based connectivity. The difference in wiring cost between and eyes closed and eyes open is maximum in the alpha frequency band when the functional connectivity is calculated using the phase-based connectivity, as the alpha desynchronization hypothesis would predict.

Figure 4 shows that the wiring cost difference, wiring cost in the eyes closed condition minus wiring cost in eyes open condition, calculated using phase-based connectivity (ISPC and PLI) is positive in the alpha band (highlighted in green) and maximum compared with the other frequency bands (delta, theta, low and high beta, and gamma). This is in agreement with the alpha desynchronization hypothesis, which stipulates that there is a loss of connections with the transition from eyes closed to eyes open. The wiring cost difference calculated using phase-lag index (PLI) is always negative except in the alpha band, that is, the wiring cost is greater for eyes open than for eyes closed. For phase-based connectivity using ISPC, the difference between wiring cost in eyes closed and eyes open conditions is positive in all bands except for delta and alpha bands. The wiring cost difference for power-based connectivity is positive for delta, theta and alpha and negative for faster frequencies.

However, the results in Figure 4 are averaged for all electrode locations and the electrode implants are not identical for across participants. To address this issue we investigate the effect of a higher degree of alertness (going from eyes closed to eyes open) for specific regions of interest. The results are shown in Figure 5. The alpha desynchronization holds for hippocampal, depth, temporal and frontal electrodes (Figures 5 a-d). On the other hand, the wiring cost difference is negative, that is, eyes open is more costly than eyes closed, in interhemispheric and grid electrodes (Figures 5 e-f).

**Figure 5:**
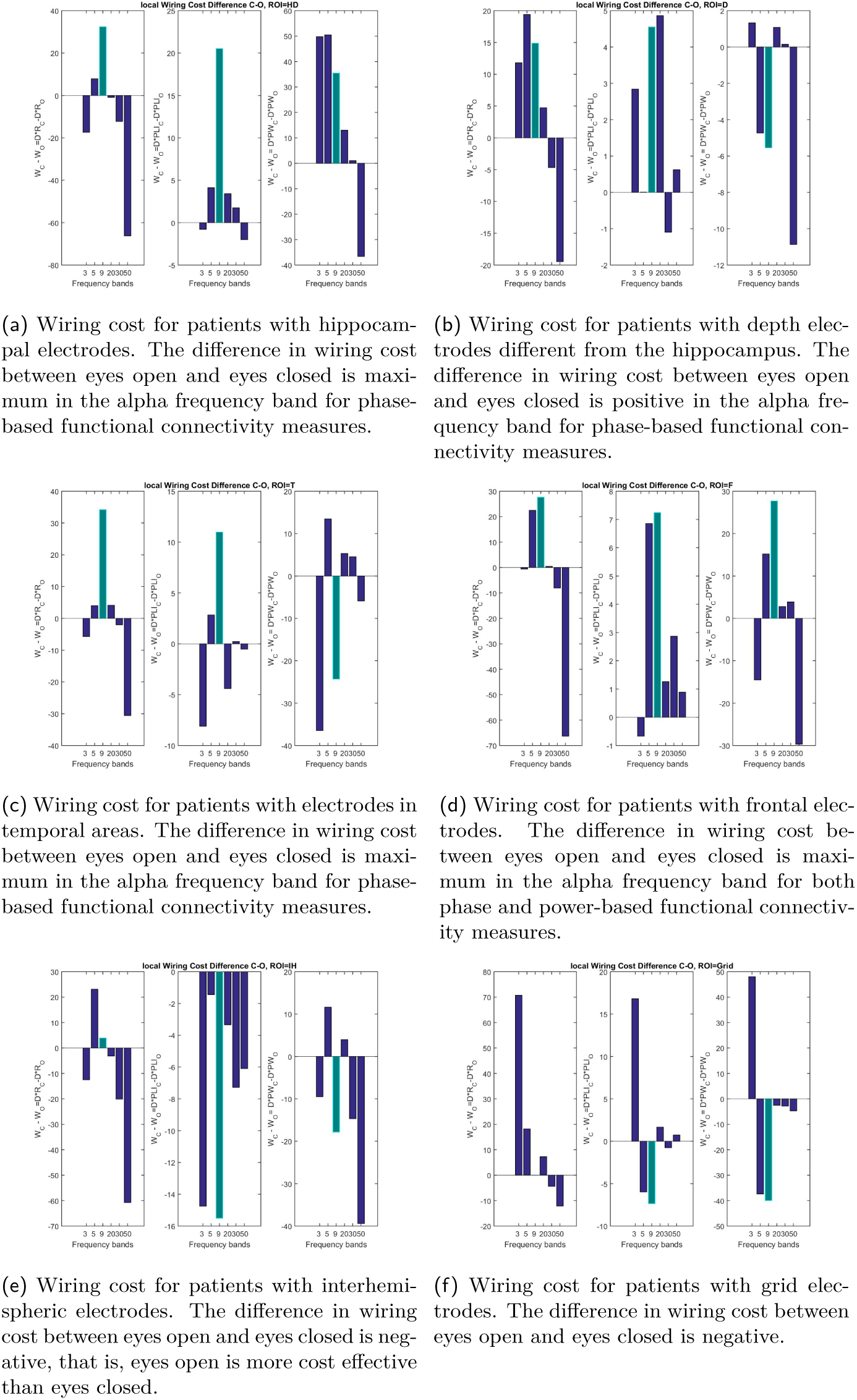
Wiring cost difference, eyes closed minus eyes open, for regions of interest. From upper-left clockwise: hippocampal, depth, temporal, frontal, inter hemispheric and grid electrodes. The wring cost is calculated as the product of the distance and the functional connectivity, from left to right, ISPC, PLI and power. In agreement with the alpha desynchronization hypothesis, the wiring cost difference is positive in the alpha frequency band (highlighted in green) in all regions except for interhemispheric (5e) and grid electrodes (5f).

### 3.2 Mesoscopic Wiring cost

Now we calculate the mesoscopic wiring cost (Equation 5). Figure 6 shows the difference between the mesoscopic wiring cost for the eyes closed and eyes open conditions, averaged across participants. The mesoscopic wiring cost calculus takes into account the connectivity pattern of the electrodes. The PLI-based (Figure 6 center) and power-based wiring cost (Figure 6 right) support the alpha desynchronization hypothesis. For ISPC-based connectivity, the wiring cost in eyes open is slightly larger than for eyes closed. Results show that the wiring cost difference (wiring cost in eyes closed minus wiring cost in eyes open), *W_C_ – W*_O_, calculated with phase-based connectivity using PLI is always negative, except for in the alpha band. That is, the wiring cost calculated using the phase-lag index is greater for eyes open than for eyes closed. For phase-based connectivity using ISPC, the difference between wiring cost between the two conditions is positive in all bands except for delta and alpha. The wiring cost difference for power-based connectivity is positive for delta, theta and alpha and negative for faster frequencies.

**Figure 6:**
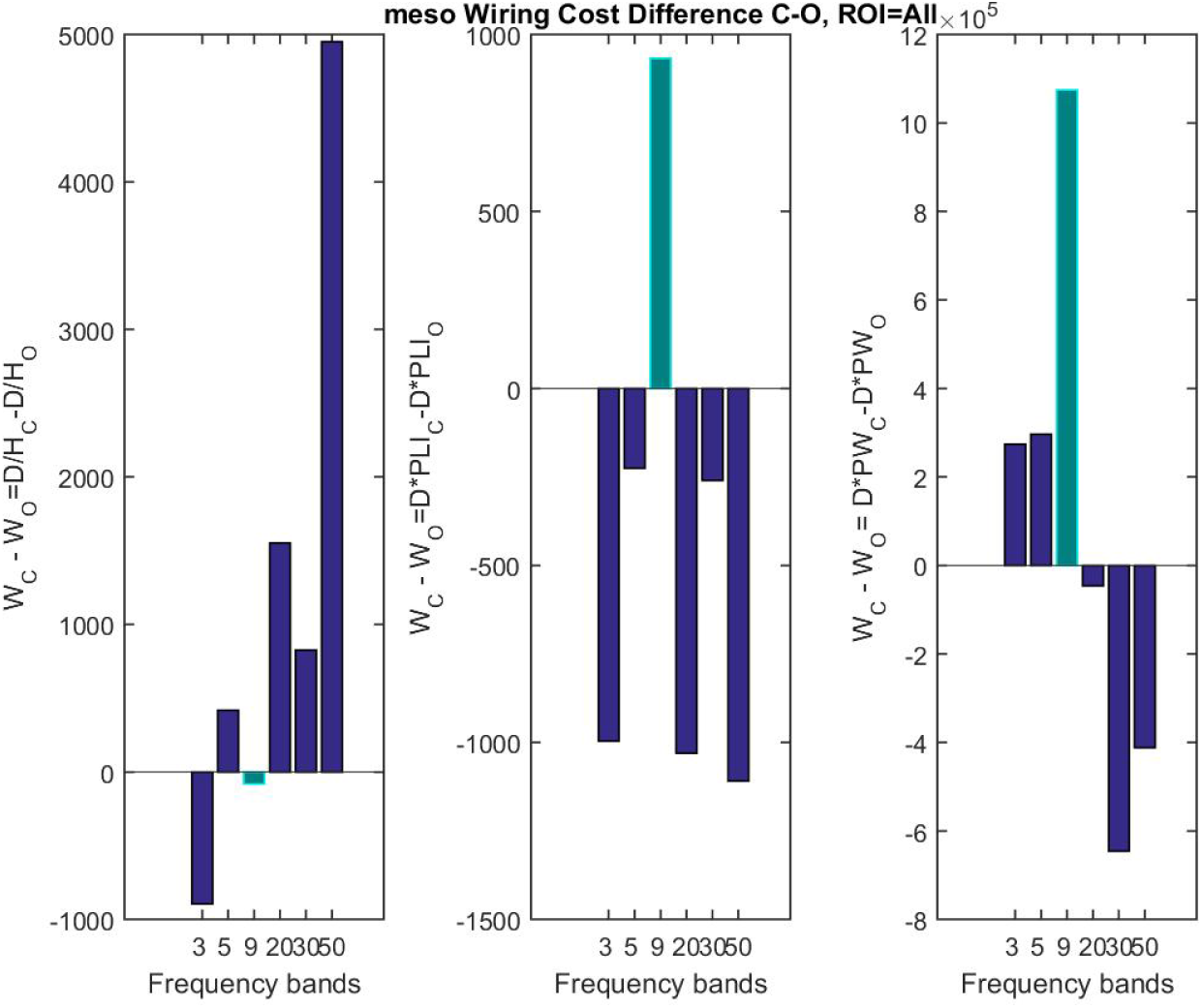
Figure shows the difference in wiring cost between eyes closed and eyes open, averaged for all patients across six frequency bands, delta, theta, alpha, low and high beta, and gamma. The wiring cost between two electrodes is calculated taking into account their connectivity patterns. From left to right, the wiring cost difference for phase-based connectivity using ISPC, PLI and power-based connectivity. The difference in wiring cost between eyes open and eyes closed is maximum in the alpha frequency band, when functional connectivity is calculated using the phase-lag index and power based measures.

PLI averages the sign of the imaginary part of the cross spectral density, thus the difference in the PLI between the two conditions, can be positive for two reasons: the average of the sign of the phases is larger in absolute value for eyes closed than for eyes open, or the absolute value of the average sign for eyes open is larger than for eyes closed and is negative. The largest positive difference of wiring cost between the conditions for PLI and power-based functional connectivity, is in the alpha band (green bar in Figure 6).

The results in Figure 6 are, however, averaged for all electrode locations and the electrode implants are not identical across participants. As was done for local wiring cost, we investigate the effect of a higher degree of alertness (going from eyes closed to eyes open) for the various regions of interest. The results are shown in Figure 7. Contrary to what was observed with local wiring cost, in mesoscopic wiring cost, the interhemispheric (Figure 6-e) and grid electrodes (Figure 6-f) have a positive wiring cost difference between eyes closed and eyes open. The reverse is seen in hippocampal (Figure 6-a), depth electrodes (Figure 6-b) and frontal regions (Figure 6-d), in which the transition from eyes closed to eyes open causes an increase in the wiring cost in the alpha frequency band. This is in agreement with EEG studies that show a decrease in alpha activity across the entire cortex in response to visual stimulation (Barry et al., 2007).

**Figure 7:**
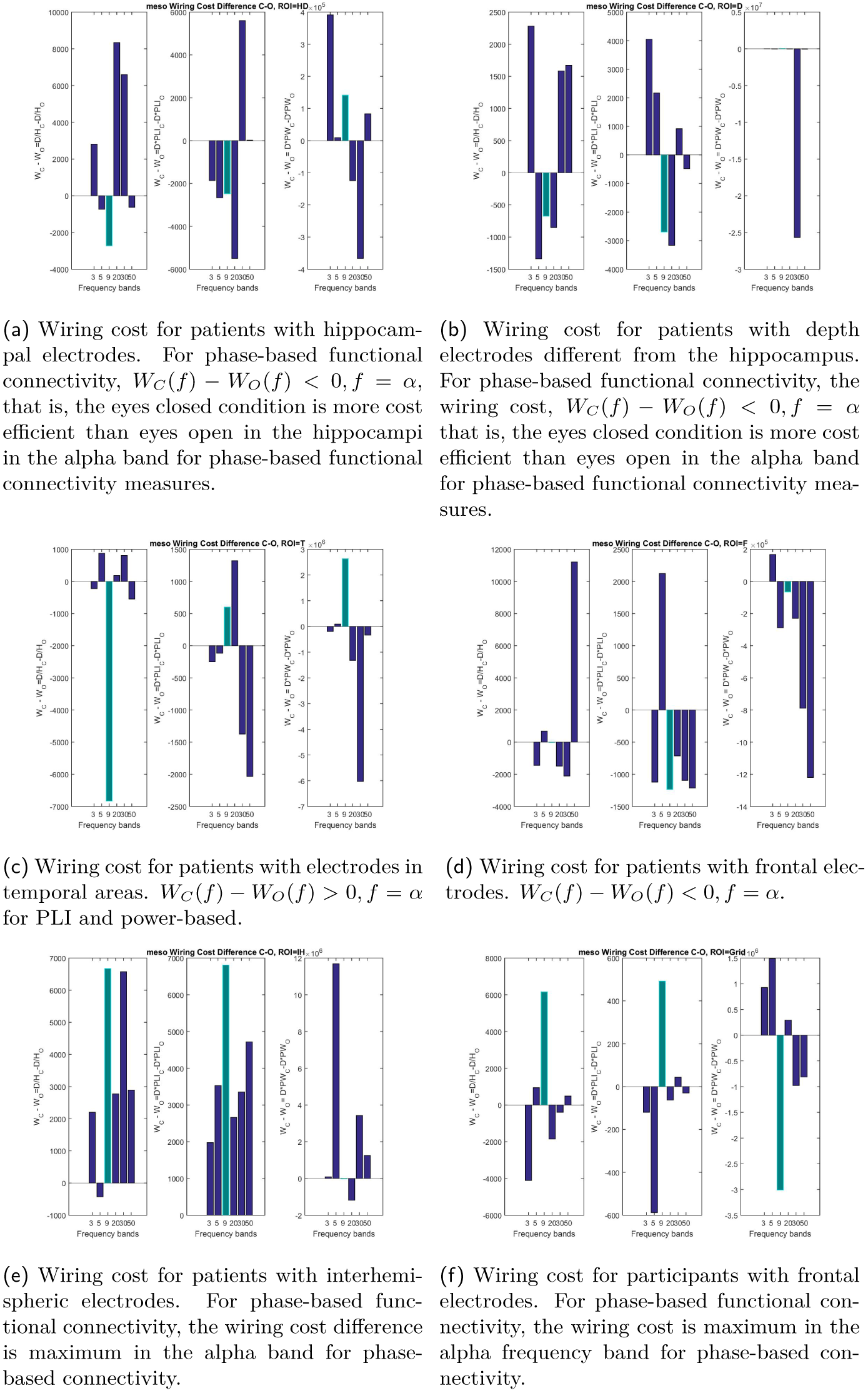
The figure shows the wiring cost difference, eyes closed minus eyes open, for regions of interest. From upper-left clockwise: hippocampal, depth, temporal, frontal, inter hemispheric and grid electrodes. The wiring cost difference is positive and maximum in the alpha frequency for inter hemispheric (7e) and grid electrodes (7f). For hippocampal (7a) and depth electrodes (7b), the opposite occurs, eyes closed is more wiring cost efficient than eyes open. Temporal (7c) electrodes show a very strong negative wiring cost difference using ISPC in the alpha band. This is due to a extreme value in one of the participants (here not shown).

Figure 4 shows that the wiring cost (Equation 4) for phase-based connectivity decreases from eyes closed to eyes open in the alpha band as the alpha desynchronization hypothesis would predict. When the wiring cost is not calculated pairwise, but also takes into account the pattern of connections between the electrodes (Equation 5) the wiring cost difference per area is reversed. While for local wiring cost, the wiring cost between conditions decreases in hippocampal, depth, temporal and frontal regions and increases in interhemispheric and grid areas, for the mesoscopic wiring cost, the opposite pattern occurs. Grid and interhemispheric electrode implants are likely subjected to a heavier informational flow than the other regions. A grid occupies a larger extension of the brain than any other implants, and interhemispheric electrodes have the longest strips. To investigate whether the alpha desynchronization hypothesis holds, we perform multivariate analysis for the mesoscopic wiring cost connectivity matrices.

### 3.3 Network analysis with perfusion method

To study the topological properties of functional connectivity networks we need to consider a threshold, which once applied to the connectivity matrix, will produce a binary graph from which network properties such as clustering, small world and others can be measured.

The selection of the threshold is, however, generally arbitrary, and the resulting network depends entirely upon that choice. We overcome this limitation by following a perfusion method, in which rather than having one threshold, we build a vector of thresholds, containing all possible threshold values between the two extreme (minimum and maximum connectivity values). For example, for the matrix *W*, of dimension *n* × *n* we obtain the threshold vector *T* with *n*^2^ elements bounded between the minimum and maximum of *W*, *T = [min(W), max(W*)].

A set of binary networks is then obtained by thresholding the wiring cost matrix for each possible threshold. Specifically, the binary matrix *B_τ_* for the threshold *τ* and wiring cost matrix *W* is such that *B_τ_* (*ij*) *=* 0 if the wiring cost between electrodes *i,j* is less than the threshold, otherwise *W*(*ij*) *< τ* and *B_τ_(ij) =* 1. Thus, for each threshold value *τ ∈ T*, we obtain a binary network and the resulting set of networks is comprised at the two extremes of the spectrum, the disconnected graph *B_τ_* (V,∅) produced when applying the threshold *τ = min(W)* and the full graph *B_τ_*(*V, E*(*W*)) resulting from applying the threshold *τ = max*(*W)*. Importantly, the set of binary networks has an internal structure that progressively increases until it becomes a fully connected network. This method is akin to perfusion in computational topology (Dabaghian et al., 2014); (Dotko et al., 2016). Figure 8 shows an example of binary networks produced for a given threshold, at the beginning of the perfusion process (8a) and halfway (8b).

**Figure 8:**
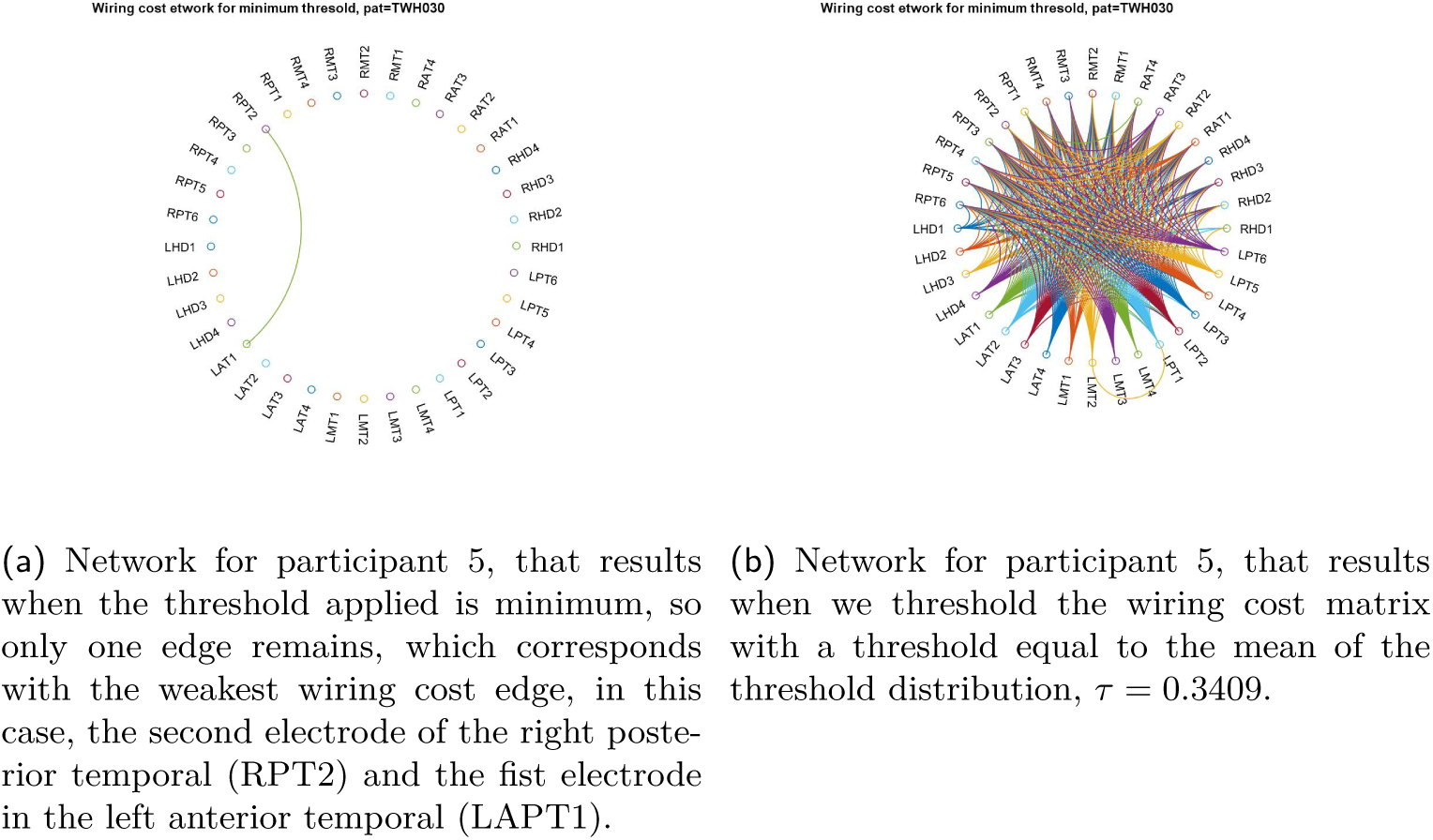
Networks that result when we apply one threshold or another, on the left the network when the minimum threshold is applied and on the right the resulting network for a threshold equal to the mean value of the wiring cost distribution.

Figure 9 shows the wiring cost difference between eyes closed and eyes open for each network derived from applying the threshold to the wiring cost matrix for two subjects. It also shows that the wiring cost difference changes depending on the network for which the wiring cost is calculated.

**Figure 9:**
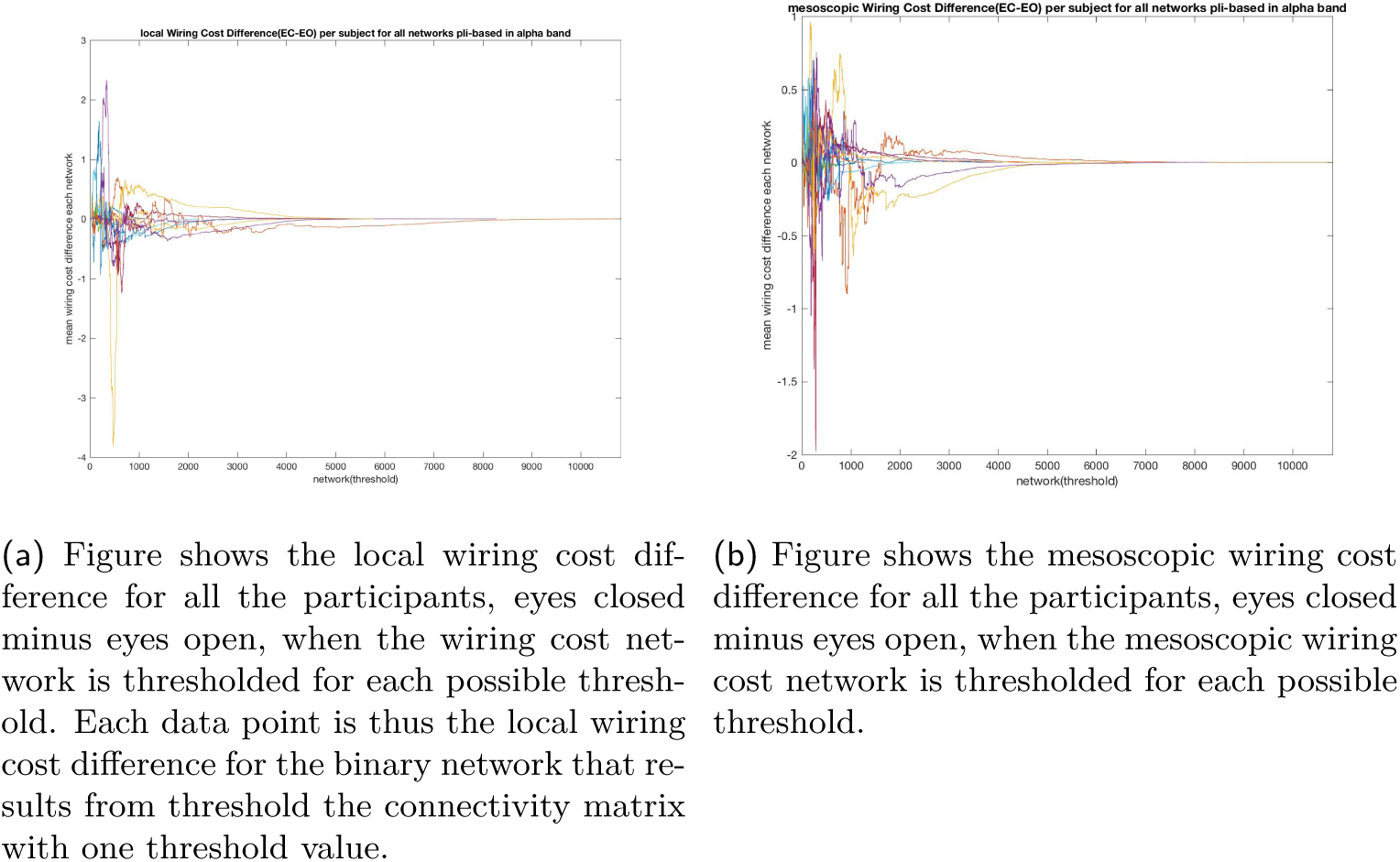
The local (9a) and mesoscopic (9b) wiring cost difference for all the participants. Each data point in the y-axis is the wiring cost difference, calculated for phase lag index in the alpha band for each network obtained by applying the threshold represented in the x-axis. The wiring cost difference oscillates between positive (the eyes closed condition has a greater wiring cost) and negative values (eyes open is more costly) to converge at 0, when the threshold approaches the maximum, and the graph is fully connected or complete (there is an edge for every pair of nodes) for both conditions.

To acquire a qualitative understanding of both conditions, eyes closed and eyes open, in terms of the wiring cost, we need to do statistics with the distribution of binary networks. The null hypothesis is that the effect of eyes closed is indistinguishable from the effect of eyes open for the wiring cost. We perform a Kruskal and Wallis test corrected for multiple comparisons. The mean ranks of the two groups, eyes closed and eyes open, are significantly different for both the local *(p >* χ̃^2^ = 4.973^−63^) and the mesoscopic wiring cost *(p > χ*̃^2^ *=* 1.65^−l8^), computed in the alpha band.

## 4 Discussion

The brain is energy hungry, it amounts to only the 2% of the weight of the body, but takes up to 20% of the body’s metabolic demand.

Yet, as with all physical systems, the brain has energy limitations. Ramón y Cajal was the first to postulate the laws of conservation of pace and material. It follows that there is a strong pressure for an efficient use of resources, for example the minimization of the synaptic wiring cost at axonal, dendritic and synaptic between nerve cells. Longer connections, and those with greater cross-sectional area, are more costly because they occupy more physical space, require greater material resources, and consume more energy per connection. Networks that strictly conserve material and space (e.g. lattice) will likely pay a price in terms of conservation of time: it will take longer to communicate an electrophysiological signal between nodes separated by the longer path lengths that are characteristic of lattices (Fornito et al., 2016). There are trade-offs between biological cost and topological value.

Functional connectivity analysis from EEG data provides an explanation for alpha desynchronization in terms of the number of connections i.e., the number of connections decreases when one’s eyes are open compared to closed. It is worth noting that the term desynchronization is defined in the literature quite vaguely, and used to mean very different things. Synchronization some times refers an to increase in band power in some frequency band (e.g. alpha) and conversely, desynchronization is also associated with a loss of power in the frequency band of interest. Stam et al.(Stam et al., 1993)provide an alternative approach to desynchronization of the alpha rhythm, which is characterized as an increase in the irregularity of the EEG signal. The EEG irregularity is quantified with the acceleration spectrum entropy (ASE), which is the normalized information entropy of the amplitude spectrum of the second derivative of a time series.

This study investigates the electrophysiological signature that characterize eyes closed and eyes open resting states in patients diagnosed with mesial lobe epilepsy, taking advantage of the unmatched spatio-temporal properties of iEEG. Both power and phase based connectivity analysis was performed for the two conditions, eyes closed and eyes open, for 6 frequency bands. Our particular focus was in the alpha band, to investigate the alpha desynchronization hypothesis. Alpha desynchronization or the alpha blocking response to eye opening was originally reported by Berger in 1929. Alpha suppression is produced by an influx of light, other afferent stimuli and mental activities (Schomer and Da Silva, 2012). Alpha rhythm is the EEG correlate of relaxed wakefulness, best obtained while the eyes are closed (Niedermeyer and da Silva, 2005).

The wiring cost, as defined here, combines the physical distance between electrodes and the statistical correlation and takes ful advantage of the spatial resolution of the ECoG signal. Specifically, the local wiring cost of two electrodes represents the product between the distance and the correlation value, while the mesoscopic wiring cost also takes the connectivity pattern of the electrodes into account to calculate their wiring cost. The combination of functional connectivity and distance networks allows us to quantify the wiring cost for the two conditions under study-eyes closed and eyes open. The rationale behind this approach is that the wiring cost might explain, at least in energy minimization terms, why, among all possible configurations, some functional connectivity patterns are selected rather than others. We mathematically define the wiring cost for a given connectivity pattern, defined in Equation 4 and Equation 5.

We find that alpha desynchronization can be explained with the wiring cost variation between eyes closed and eyes open. The local wiring cost decreases as the alpha desynchronization hypothesis predicts when going from eyes closed to eyes open. This is observed for phase based connectivity wiring cost overall and consistently for most regional electrode implants (hippocampal, depth, temporal and frontal). The mesoscopic wiring cost, however, presents more variability and we observe that the wiring cost may increase for eyes open depending on where the electrodes are located. For phase lag index based mesoscopic wiring cost, the alpha desynchronization holds in frontal and interhemispheric but not in hippocampal, depth and temporal areas.

To investigate whether the wiring cost captures the loss of connectivity predicted by the alpha desynchronization hypothesis, we calculated the distribution of wiring cost values associated with the connectivity matrix for every possible threshold value from a threshold vector bounded by the minimum and maximum functional connectivity value. We find that both the local wiring cost and the mesoscopic wiring cost defined here, decreases on average when going from eyes closed to eyes open. Furthermore, we perform network perfusion analysis to study the wiring cost distribution from multiple multiple thresholds, rather than selecting one threshold. Showing that the wiring cost in eyes closed and open resting states is statistically different.

Although intracranial electroencephalography has unmatched spatial and temporal specificity, may not the optimal method for studying macroscopic aspects of the human brain. This study has the limitation that the electrode implants tend to be located in the seizure sensitive temporal lobe and leave untouched occipital and parietal lobes. Furthermore, the participants are patients with drug resistant epilepsy, who likely have the functioning in some, if not all the brain areas where they have the implants, seriously compromised by the pathology over the years. A more straightforward model system for the study of the wiring cost difference between two connectivity patterns would be EEG or fMRI, in which the signal source is regularized in a common brain volume template.

The wiring cost metric here defined can be applied to resting state (sleep, awake), task-based and pathological conditions, for example in epileptic seizures. In a forthcoming study, we show that the wiring cost-local and mesoscopic-increases dramatically in the ictal period compared to the pre-ictal period.

This work is a step forward in understanding the electrophysiological differences between the eyes open and eyes closed resting state conditions. It uses a straight forward and easily replicable approach to investigate the electrophysiology of baseline condition in terms of energy efficiency. The minimization of the wiring cost for functional connectivity networks acting over networks of intracranial electrodes provides a new avenue to understand the electrophysiology of resting state.

## Acknowledgements

We acknowledge the support of the Bial Foundation, grant number #20614.

## Footnotes

1 intracraneal electroencephalography, iEEG, and electrocorticography, ECoG, are here used indistinctly.

2 2ISPC represents the clustering in polar space of phase angle differences between electrodes resulting from the convolution between a complex wavelet and the signal and is also referred in the literature as R (Cohen, 2014)

## References

Barry, R. J., Clarke, A. R., Johnstone, S. J., Magee, C. A., and Rushby, J. A. (2007). Eeg differences between eyes-closed and eyes-open resting conditions. Clinical Neurophysiology, 118(12):2765–2773.

Biswal, B., Yetkin, F. Z., Haughton, V. M., and Hyde, J. S. (1995). Functional connectivity in the motor cortex of resting human brain using echo-planar MRI. Magnetic resonance in medicine: official journal of the Society of Magnetic Resonance in Medicine / Society of Magnetic Resonance in Medicine, 34(4):537–541. PMID: 8524021.

Buckner, R. L. and Vincent, J. L. (2007). Unrest at rest: default activity and spontaneous network correlations. Neuroimage, 37(4):1091–1096.

Cohen, M. X. (2014). Analyzing neural time series data: theory and practice. MIT Press.

Dabaghian, Y., Brandt, V. L., and Frank, L. M. (2014). Reconceiving the hippocampal map as a topological template. Elife, 3:e03476.

Dotko, P., Hess, K., Levi, R., Nolte, M., Reimann, M., Scolamiero, M., Turner, K., Muller, E., and Markram, H. (2016). Topological analysis of the connectome of digital reconstructions of neural microcircuits. arXiv preprint arXiv:1601.01580.

Fornito, A., Zalesky, A., and Bullmore, E. (2016). Fundamentals of Brain Network Analysis. Academic Press.

Freeman, W. J. and Zhai, J. (2009). Simulated power spectral density (psd) of background electrocorticogram (ecog). Cognitive neurodynamics, 3(1):97–103.

Fukushima, M., Chao, Z. C., and Fujii, N. (2015). Studying brain functions with mesoscopic measurements: Advances in electrocorticography for non-human primates. Current opinion in neurobiology, 32:124–131.

Geller, A. S., Burke, J. F., Sperling, M. R., Sharan, A.D., Litt, B., Baltuch, G. H., Lucas, T. H., and Kahana, M. J. (2014). Eye closure causes widespread low-frequency power increase and focal gamma attenuation in the human electrocorticogram. Clinical Neurophysiology., 125(9):1764–1773.

Greicius, M. D. and Menon, V. (2004). Default-mode activity during a passive sensory task: uncoupled from deactivation but impacting activation. Journal of cognitive neuroscience, 16(9):1484–1492.

Llinás, R. R. (1988). The intrinsic electrophysiological properties of mammalian neurons: insights into central nervous system function. Science (New York, N.Y.), 242(4886):1654–1664.

Maandag, N. J., Coman, D., Sanganahalli, B. G., Herman, P., Smith, A. J., Blumenfeld, H., Shulman, R. G., and Hyder, F. (2007). Energetics of neuronal signaling and fmri activity. Proceedings of the National Academy of Sciences, 104(51):20546–20551.

Mantini, D., Perrucci, M. G., Del Gratta, C., Romani, G. L., and Corbetta, M. (2007). Electrophysiological signatures of resting state networks in the human brain. Proceedings of the National Academy of Sciences, 104(32):13170–13175.

Morcom, A. M. and Fletcher, P. C. (2007). Does the brain have a baseline? why we should be resisting a rest. Neuroimage, 37(4):1073–1082.

Mormann, F., Lehnertz, K., David, P., and Elger, C. E. (2000). Mean phase coherence as a measure for phase synchronization and its application to the eeg of epilepsy patients. Physica D: Nonlinear Phenomena, 144(3):358–369.

Musso, F., Brinkmeyer, J., Mobascher, A., Warbrick, T., and Winterer, G. (2010). Spontaneous brain activity and eeg microstates. a novel eeg/fmri analysis approach to explore resting-state networks. Neuroimage, 52(4):1149–1161.

Niedermeyer, E. and da Silva, F. L. (2005). Electroencephalography: basic principles, clinical applications, and related fields. Lippincott Williams & Wilkins.

Nolte, G., Bai, O., Wheaton, L., Mari, Z., Vorbach, S., and Hallett, M. (2004). Identifying true brain interaction from eeg data using the imaginary part of coherency. Clinical neurophysiology, 115(10):2292–2307.

Nolte, G., Ziehe, A., Nikulin, V. V., Schlögl, A., Krämer, N., Brismar, T., and Müller, K.-R. (2008). Robustly estimating the flow direction of information in complex physical systems. Physical review letters, 100(23):234101.

Northoff, G., Duncan, N. W., and Hayes, D. J. (2010). The brain and its resting state activity? experimental and methodological implications. Progress in neurobiology, 92(4):593–600.

Papo, D. (2013). Why should cognitive neuroscientists study the brain’s resting state? Frontiers in human neuroscience, 7:45.

Patriat, R., Molloy, E. K., Meier, T. B., Kirk, G. R., Nair, V. A., Meyerand, M. E., Prabhakaran, V., and Birn, R. M. (2013). The effect of resting condition on resting-state fmri reliability and consistency: a comparison between resting with eyes open, closed, and fixated. Neuroimage, 78:463–473.

Peraza, L. R., Asghar, A. U., Green, G., and Halliday, D. M. (2012). Volume conduction effects in brain network inference from electroencephalographic recordings using phase lag index. Journal of neuroscience methods, 207(2):189–199.

Schneider, F., Bermpohl, F., Heinzel, A., Rotte, M., Walter, M., Tempelmann, C., Wiebking, C., Dobrowolny, H., Heinze, H., and Northoff, G. (2008). The resting brain and our self: self-relatedness modulates resting state neural activity in cortical midline structures. Neuroscience, 157(1):120–131.

Schomer, D. L. and Da Silva, F. L. (2012). Niedermeyer’s electroencephalography: basic principles, clinical applications, and related fields. Lippincott Williams & Wilkins.

Sokoloff, L., Mangold, R., Wechsler, R. L., Kennedy, C., and Kety, S. S. (1955). The effect of mental arithmetic on cerebral circulation and metabolism. Journal of Clinical Investigation, 34(7 Pt 1):1101.

Stam, C., Tavy, D., and Keunen, R. (1993). Quantification of alpha rhythm desynchronization using the acceleration spectrum entropy of the eeg. Clinical EEG and Neuroscience, 24(3):104–109.

Stam, C. J., Nolte, G., and Daffertshofer, A. (2007). Phase lag index: assessment of functional connectivity from multi channel eeg and meg with diminished bias from common sources. Human brain mapping, 28(11):1178–1193.

Tan, B., Kong, X., Yang, P., Jin, Z., and Li, L. (2013). The difference of brain functional connectivity between eyes-closed and eyes-open using graph theoretical analysis. Computational and mathematical methods in medicine, 2013.

Tracy, J. I. and Doucet, G. E. (2015). Resting-state functional connectivity in epilepsy: growing relevance for clinical decision making. Current opinion in neurology, 28(2):158–165.

Van Den Heuvel, M. P. and Pol, H. E. H. (2010). Exploring the brain network: a review on resting-state fmri functional connectivity. European Neuropsychopharmacology, 20(8):519–534.

Vinck, M., Oostenveld, R., van Wingerden, M., Battaglia, F., and Pennartz, C. M. (2011). An improved index of phase-synchronization for electrophysiological data in the presence of volume-conduction, noise and sample-size bias. Neuroimage, 55(4):1548–1565.

Wang, L., Zang, Y., He, Y., Liang, M., Zhang, X., Tian, L., Wu, T., Jiang, T., and Li, K. (2006). Changes in hippocampal connectivity in the early stages of alzheimer’s disease: evidence from resting state fmri. Neuroimage, 31(2):496–504.

Yan, C., Liu, D., He, Y., Zou, Q., Zhu, C., Zuo, X., Long, X., and Zang, Y. (2009). Spontaneous brain activity in the default mode network is sensitive to different resting-state conditions with limited cognitive load. PloS one, 4(5):e5743.

